# Terminal type-specific cannabinoid CB1 receptor alterations in patients with schizophrenia: a pilot study

**DOI:** 10.1101/2023.04.11.536217

**Authors:** Shinnyi Chou, Kenneth N Fish, David A Lewis, Robert A Sweet

## Abstract

**Background:** Individuals with schizophrenia are at elevated genetic risks for comorbid cannabis use, and often experience exacerbations of cognitive and psychotic symptoms when exposed to cannabis. These findings have led a number of investigators to examine cannabinoid CB1 receptor (CB1R) alterations in schizophrenia, though with conflicting results. We recently demonstrated the presence of CB1R in both excitatory and inhibitory boutons in the human prefrontal cortex, with differential levels of the receptor between bouton types. We hypothesized that the differential enrichment of CB1R between bouton types – a factor previously unaccounted for when examining CB1R changes in schizophrenia – may resolve prior discrepant reports and increase our insight into the effects of CB1R alterations on the pathophysiology of schizophrenia.

**Methods:** Using co-labeling immunohistochemistry and fluorescent microscopy, we examined total CB1R levels and CB1R levels within excitatory (vGlut1-positive) and inhibitory (vGAT-positive) boutons of prefrontal cortex samples from ten pairs of individuals diagnosed with schizophrenia and non-psychiatric comparisons.

**Results:** Significantly higher total CB1R levels were found within samples from individuals with schizophrenia. Terminal type-specific analyses identified significantly higher CB1R levels within excitatory boutons in samples from individuals with schizophrenia relative to comparisons. In contrast, CB1R levels within the subset of inhibitory boutons that normally express high CB1R levels (presumptive cholecystokinin neuron boutons) were lower in samples from individuals with schizophrenia relative to comparison samples.

**Conclusion:** Given CB1R’s role in suppressing neurotransmission upon activation, these results suggest an overall shift in excitatory and inhibitory balance regulation toward a net reduction of excitatory activity in schizophrenia.

## Introduction

Cannabis is the most widely used recreational psychoactive substance worldwide, with ongoing increase in usage(1,2). Cannabis use is associated with various psychiatric comorbidities(3) and represents one of the strongest environmental factors associated with schizophrenia (SZ)(4). In population studies, cannabis use diagnoses significantly increase the hazard ratios of developing SZ(5,6). Temporally, cannabis use is associated with younger ages of first psychotic episodes(7), with severity of cannabis use dose-dependently modulating the risk of developing SZ(8,10), and many patients with cannabis-induced psychosis later develop SZ(9). Importantly, individuals diagnosed with SZ use cannabis at significantly higher rates, with exposure to cannabis transiently exacerbating cognitive and positive symptoms(11), and a history of cannabis use being associated with worse illness prognosis(12,13).

Recent genetic studies investigating the relationship between cannabis use and SZ suggest that shared risk liabilities of cannabis use disorder and SZ may partially account for these observations. In particular, Mendelian randomization studies demonstrated that while SZ increases the risk of cannabis use(14), cannabis use further increases the risk of SZ beyond what can be accounted for by genetic correlations alone(15). These findings suggest that changes within the endocannabinoid system may affect the clinical outcomes of both cannabis use and SZ, and increasing our understanding of this system remains critical, with potential therapeutic benefits.

Δ9-tetrahydrocannabinol (THC), the major psychoactive substance in cannabis, targets the cannabinoid CB1 receptor (CB1R)(16). A ubiquitous G-protein coupled receptor (GPCR)(17), CB1R is highly expressed within the cholecystokinin (CCK) subtype GABAergic inhibitory interneurons within the human cortex (18,19). However, it is also found in other interneuronal subtypes and glutamatergic neurons(20,21). Functionally, presynaptic CB1R activation results in reduced synaptic transmission by attenuating neurotransmitter release, a phenomenon known as depolarization-induced suppression of inhibition (DSI) at inhibitory boutons, and depolarization-induced suppression of excitation (DSE) at excitatory sites(22–24). Thus, CB1R appears to be critical in regulating cortical excitatory-inhibitory (E/I) balance(25).

CB1R alterations may also play a role in cognitive impairments in SZ – a function involving the prefrontal cortex (PFC)(26). Studies have identified reduced dendritic spine density in PFC pyramidal neurons of individuals with SZ(27–29), which – given the spines’ role in forming excitatory synapses – may suggest decreased excitatory drive(30,31). In contrast, reduced mRNA expression of gamma-aminobutyric acid (GABA) synthesizing enzymes in SZ suggests decreased GABA synthesis(32,33), which may also contribute to E/I balance impairments. However, the full circuitry abnormalities leading to these PFC disturbances in SZ remain uncertain.

Considering the above, studies have investigated CB1R changes as potential mechanisms in the pathophysiology of SZ(34,35). Interestingly, ligand-binding autoradiography studies targeting all CB1R, including those in non-CCK interneurons and glutamatergic neurons, demonstrated increased cortical CB1R binding in postmortem samples from individuals with SZ(36–40). In contrast, immunohistochemistry (IHC) antibody-based studies showed decreased CB1R protein levels in SZ(41–43). Of note, existing literature indicates that the antibodies used in prior IHC studies preferentially labeled puncta with high CB1R expressions in inhibitory boutons – predominantly found to be CCK-positive cells given the high abundance and expression of CB1R within CCK subtype GABAergic interneurons(40,43,44). We hypothesized that the discrepant results between these two methods may stem from Terminal type-specific alterations of CB1R levels in SZ. We undertook to preliminarily test this idea by performing quantitative IHC using a CB1R antibody we previously showed to detect CB1R in both interneurons and glutamatergic neurons in the postmortem human PFC(21), in an existing cohort of SZ subjects previously examined with both ligand-binding autoradiography and CCK cell selective CB1R antibodies.

## Methods and materials

### Human Tissue

We studied ten individuals with SZ, each matched for sex and age to an unaffected comparison subject (Ctrl) without psychiatric diagnoses (Table 1 and S1). To control for experimental variance, subjects from each pair were processed together throughout the protocol. All pairs were previously assayed for PFC CB1R levels using both ligand-binding and antibody-based approaches(40,43). See Table 2 for each pair’s ligand-binding and antibody-based CB1R ratios.

**Table 1.** Summary characteristics of individuals included in the study.

| **Characteristics** | **Ctrl, N = 10** | **SZ, N = 10** | **p-value** |
| --- | --- | --- | --- |
| Age, years, mean (SD) | 46.9 (15.9) | 48.4 (13.7) | 0.82 |
| Sex, n |  |  |  |
| Female | 1 | 1 | 1.00 |
| Male | 9 | 9 |  |
| Race, n |  |  |  |
| Black | 2 | 1 | 1.00 |
| White | 8 | 9 |  |
| Cannabis use history |  |  | 0.21 |
| Yes | 0 | 3 |  |
| No | 10 | 7 |  |
| PMI, hours, mean (SD) | 17.9 (5.9) | 20.3 (11.2) | 0.55 |
| Storage time, months, mean (SD) | 187.8 (16.3) | 188.8 (24.4) | 0.92 |
| pH, mean (SD) | 6.96 (0.24) | 6.92 (0.20) | 0.69 |
| Leading cause of death (% subjects affected) | Cardiovascular (70%) | Cardiovascular (60%) |  |

Abbreviations: Ctrl = unaffected comparison, SZ = schizophrenia, PMI = postmortem interval, SD = standard deviations

**Table 2.** Reciprocal ligand-binding and IHC protein ratio results obtained from subject pairs used in prior studies. These pairs are included in the current study (i.e., those with demographic information provided in Table 1). IHC protein ratio is calculated as the percentage of protein level (measured in optical density) in samples from subjects with schizophrenia (SZ) to samples from unaffected comparisons. Ligand-binding ratio is calculated as the percentage of OMAR ligand-binding (fm/mg) in samples from patients with SZ to samples from unaffected comparisons. Magnitude difference is calculated as the difference between the ligand binding ratio and the protein IHC ratio.

| **Pair** | **IHC protein**<br>**SZ/Ctrl ratio (%)** | **Ligand-binding**<br>**SZ/Ctrl ratio (%)** | **Magnitude difference**<br>**(Ligand-binding - protein)** |
| --- | --- | --- | --- |
| 1 | -5.57 | 25.76 | 31.33 |
| 2 | -11.04 | 41.18 | 52.22 |
| 3 | -20.18 | 2.51 | 22.68 |
| 4 | -31.61 | 74.85 | 106.46 |
| 5 | -24.01 | -0.74 | 23.27 |
| 6 | -6.17 | 29.19 | 35.35 |
| 7 | -30.19 | 43.08 | 73.27 |
| 8 | -23.15 | -5.18 | 17.97 |
| 9 | -14.64 | 47.74 | 62.38 |
| 10 | -21.75 | 14.52 | 36.27 |
| Mean (SD) | -18.83 (9.21) | 27.29 (25.27) | 46.12 (27.84) |

Abbreviations: Ctrl = unaffected comparison, SZ = schizophrenia, IHC = immunohistochemistry

Brain specimens from subjects were obtained from autopsies conducted at the Allegheny County Office of the Medical Examiner, Pittsburgh, PA, following consent for donation from next of kin. Psychiatric or neurological histories were determined by an independent committee of experienced research clinicians using information obtained from clinical records and structured interviews conducted with a surviving relative, including any known history of cannabis use or use disorders. The University of Pittsburgh’s Committee for the Oversight of Research and Clinical Trials Involving Decedents and Institutional Review Board for Biomedical Research approved all procedures.

Following brain retrieval, left hemispheres were cut into 1.0-2.0cm-thick coronal blocks, fixed for 48h in phosphate-buffered 4% paraformaldehyde at 4°C, immersed in graded cold sucrose solutions, and stored at -30°C in cryoprotectant solutions until sectioning(45). PFC blocks containing the region of interest (ROI; Brodmann area 9) were sectioned coronally at 40µm on a cryostat, and every 40th section was Nissl stained to serve as anatomical references for laminar identification. Unstained sections were stored in cryoprotectant solution at -30°C until processed for immunohistochemistry.

### Immunohistochemistry

One free-floating tissue section per subject containing the ROI was used. Sections were washed in 0.1M phosphate-buffered saline (PBS) then incubated for 75min in 0.01M sodium citrate solution at 80°C to retrieve antigens(46). After cooling to room temperature (RT), sections were immersed in 1% sodium borohydride for 30min at RT to reduce background autofluorescence(47), followed by membrane permeabilization with 0.3% Triton X-100 in PBS for 30min at RT. Sections were blocked with 20% normal goat serum (NGS) in PBS for 2h at RT to reduce nonspecific antibody binding, then incubated for 72h at 4°C in PBS containing 2% NGS and primary antibodies.

Primary antibodies included monoclonal mouse anti-vGAT antibody (1:500; Synaptic Systems, Göttingen, Germany; product # 131011) – which labels inhibitory boutons; polyclonal guinea pig anti-vGlut1 antibody (1:500; Millipore Sigma, Burlington, MA; product # AB5905) – which labels excitatory boutons, and polyclonal rabbit anti-CB1R antibody (1:2000; Synaptic Systems, Göttingen, Germany; product # 258003). We previously demonstrated successful and specific vGAT and vGlut1 labeling in human and non-human primate postmortem studies using these antibodies(48–51). The CB1R antibody demonstrated successful co-labeling with both vGAT and vGlut1 in both neuronal cultures and postmortem human brain samples(21,52). In addition, vGAT and CB1R antibody specificities were validated through knockout samples(53,54), and vGlut1 antibody through pre-adsorption controls (Millipore certificate of analysis, 2016).

Post primary antibody incubation, sections were rinsed for 4×30min in PBS and incubated for 24h at 4°C in PBS containing 2% NGS and goat host secondary antibodies conjugated to Alexa-488 (1:500; vGlut1), Alexa-568 (1:500; CB1R) and Alexa-647 (1:500; vGAT; Invitrogen, Grand Island, NY, for Alexa antibodies). Sections were rinsed for 4×30min in PBS, mounted on slides, cover slipped (ProLong Gold antifade reagent, Invitrogen), sealed with clear nail polish along coverslip edges, and stored at 4°C until imaged. A sample CB1R-immunoreactive (IR) labeling within postmortem PFC is shown in Figure 1, where CB1R-IR signals are seen co-localized with vGAT-IR and vGlut1-IR puncta. There are also CB1R-IR labeling of neuronal soma and axons not co-localized with either synaptic marker, i.e., CB1R-IR puncta that are neither vGlut-IR nor vGAT-IR.

**Figure 1.**
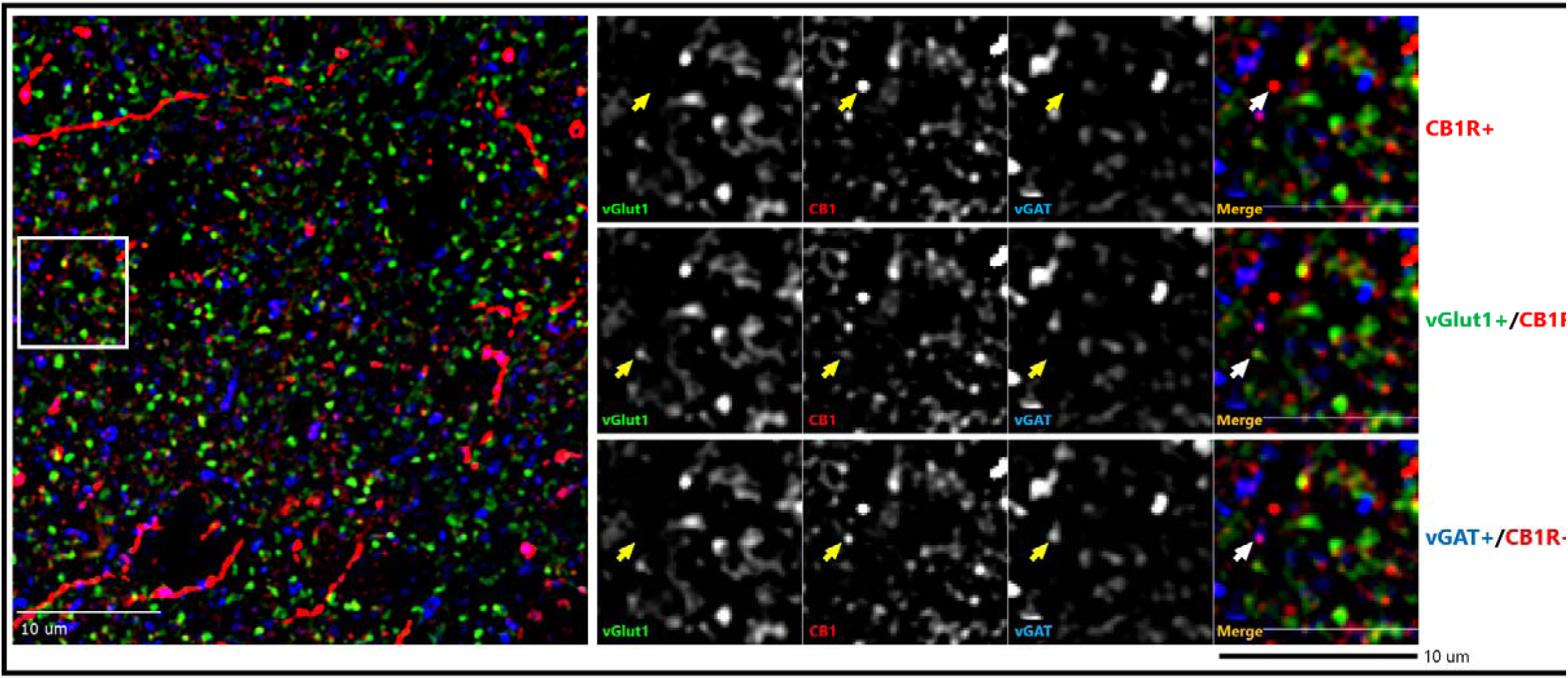
Representative micrograph of immunohistochemical labeling of postmortem human prefrontal cortex tissue section. Left panels: Puncta with vGlut1-immunoreactive (IR) (green), vGAT-IR (blue) & CB1R-IR (red) labeling are distributed throughout the image field. Right panels: Enlarged images with arrows identifying puncta with antibody immunoreactivity toward a single or multiple proteins.

### Tissue sampling

Sampling procedure was as previously described(55,56). Contours outlining each section were drawn in Stereo Investigator version 8 (MicroBrightField Inc., Natick, MA). To ensure representative sampling of complete gray matter, image stacks were obtained from six to ten randomly chosen sites for each layer per subject, determined using nearby Nissl-stained sections, equally sampled within and across subject pairs. Boundaries of cortical layers were estimated as percent of distance from pial surface to white matter: Layer 1 (pia–10%), Layer 2 (10–20%), Layer 3 (20–50%), Layer 4 (50– 60%), Layer 5 (60–80%), Layer 6 (80%–gray/white matter border)(43,57). Tissue thickness (z-axis depth) for each site was measured and divided by 40µm (original section thickness) to correct for shrinkage during IHC.

### Confocal microscopy

Microscopy equipment and capturing parameters were as previously described(58). Data were collected using a 60×1.40 numerical aperture super-corrected oil immersion objective mounted on an Olympus BX51Wl upright microscope (Olympus America Inc., Center Valley, PA) equipped with an Olympus spinning disk confocal unit, Hamamatsu Orca R2 camera (Hamamatsu, Bridgewater, NJ), MBF CX9000 front mounted digital camera (MicroBrightField Inc., Natick, MA), BioPrecision2 XYZ motorized stage with linear XYZ encoders (Ludl Electronic Products Ltd., Hawthorne, NY), excitation and emission filter wheels (Ludl Electronic Products Ltd., Hawthorne, NY), Sedat Quad 89000 filter set (Chroma Technology Corp., Bellows Falls, VT), and Lumen 220 metal halide lamp (Prior Scientific, Rockland, MA).

Equipment was controlled by SlideBook 6.0 (Intelligent Imaging Innovations, Inc., Denver, CO), which was also used for post-image processing. Three-dimensional image stacks (two-dimensional images successively captured at 0.25µm z-dimension intervals) were acquired with a depth spanning top 20% of tissue thickness (i.e., measuring 20% of thickness beginning at the coverglass), starting from the plane furthest away from the coverglass and stepping up until reaching tissue surface. Images were 512×512 pixels (55×55µm) in the XY dimension. Stacks were collected using optimal exposure settings (i.e., those yielding the greatest dynamic range possible for the camera without saturated pixels). Z-positions were normalized to original section thickness and exposures normalized for each capture post-image processing prior to analysis.

### Image processing

Images were processed as previously described(58,59), using SlideBook and Automation Anywhere software (Automation Anywhere, Inc., San Jose, CA). Image stacks were deconvolved using AutoQuant’s blind deconvolution algorithm (MediaCybernetics, Rockville, MD). After deconvolution, separate Gaussian channels were made for each deconvolved channel by calculating a difference of Gaussians (sigma 0.7 - sigma 2.0). These channels, which enhanced immunofluorescence edge demarcations, were used for data segmentation.

Segmentation of Gaussian channels was performed using a previously described iterative combined intensity/morphologic thresholding algorithm with MATLAB (MATLAB, The MathWorks Inc., Natick, MA)(48). After obtaining initial values for iterative segmentation for each channel using Otsu’s method within SlideBook, each subsequent iteration increased threshold by 50 gray levels, and object masks were size gated within 0.03–2.0µm^3^. After each segmentation, masked objects were merged with prior iterations, with final resulting masks copied back onto the original deconvolved channels (i.e., without Gaussian subtraction) to obtain pixel intensity information. Lipofuscin, an autofluorescent lysosomal degradation product, which may confound quantitative fluorescence measures in human postmortem tissues, was imaged using a separate channel at a constant exposure time across all sections.

After generating vGAT and vGlut1 bouton object masks, mean CB1R intensity in analog-to-digital units (ADU) underneath each masked object was obtained. Values were averaged across all boutons per sampled site for each bouton type. The resulting Terminal type-specific mean CB1R intensity values for each sampled site were then averaged across each layer to obtain a single value as the dependent measure. For determining all CB1R signal irrespective of terminal type within each site, sum CB1R intensity was measured from a single 2D plane, and site values averaged for each layer as the dependent measure.

Prior to analyses, data were filtered to ensure accurate representation of receptor labeling. Based upon examination of antibody signal penetrance across tissue thickness, only objects falling within 10–14µm from tissue surface after correcting for tissue shrinkage were included for analysis. To prevent potential spherical aberration confounding measurements, a virtual counting frame inclusive of signals falling between the upper and lower 2% of XY dimensions was used (i.e., between 10-502 units for each dimension). To ensure accurate capture of Terminal type-specific measurements, objects overlapping the lipofuscin and both vGlut1 and vGAT masks were excluded from analysis.

### Statistical analysis

Demographic data were analyzed using Fisher’s exact test (categorical variables) and Student’s two-tailed T-test (continuous variables). To analyze sum CB1R intensity, analysis of covariance (ANCOVA) models were performed. Sum CB1R intensity values at all sampled sites per cortical layer per subject were averaged to obtain a single measure as the dependent variable. Subject group, cortical layer, and subject group × cortical layer two-way interaction were entered as fixed effects, and subject pair entered as a blocking factor. To assess possible confounding effects of cohort variables (sex, race, age, postmortem interval, and tissue storage time), a second unpaired ANCOVA model was performed to validate the first model, using subject group, cortical layer, and subject group × cortical layer two-way interaction as fixed effects, and cohort variables as covariates. Results for both paired and unpaired models were reported.

As existing literature indicates that within GABAergic interneurons, CB1R is most abundant and expressed at highest levels within CCK-positive cells, and given that prior IHC studies used an anti-CB1R antibody that specifically labeled inhibitory boutons with high CB1R expressions determined to be predominantly CCK-positive subtype interneurons(40,43,44,60,61), we separated inhibitory bouton populations for Terminal type-specific analysis of mean CB1R intensity. Inhibitory boutons were categorized as high- or low-CB1R-expressors, using the median value of non-psychiatric subjects’ mean CB1R intensities in vGAT-IR boutons (712 ADU) to define groups after reviewing total intensity distribution (see Figure S1). We then compared mean CB1R intensities within excitatory (i.e., vGlut1-IR), high-CB1R-expressing, and low-CB1R-expressing inhibitory (i.e., vGAT-IR) boutons between groups.

To analyze mean CB1R intensity, ANCOVA models were performed. Mean CB1R intensity values for each terminal type at all sampled sites per cortical layer per subject were averaged to obtain a single measure as the dependent variable. Subject group, cortical layer, terminal type, subject group × terminal type two-way interaction, and subject group × cortical layer × terminal type three-way interaction were entered as fixed effects, and subject pair entered as a blocking factor. A second unpaired ANCOVA model was performed to validate the first model, using the same fixed effects as the first model, and cohort variables as covariates. Results for both models were reported.

To analyze effects of cannabis and medications on CB1R, independent-samples two-tailed T-test was used to compare within-pair ratios of mean or sum CB1R intensities (Ctrl/SZ CB1R intensity ratio) between pairs with and without cannabis or medication exposure histories in SZ subjects. When appropriate, significant differences were followed by *post hoc* Bonferroni tests to correct for increased risk of a type I error when making multiple comparisons. For all analyses, *p* < 0.05 was considered statistically significant.

## Results

### Global CB1R alterations in the PFC of patients with SZ

Sum CB1R intensity (encompassing vGlut1-IR, vGAT-IR, and non-vGlut1-IR/vGAT-IR populations) was significantly +26.8% higher in SZ compared to Ctrl, F(1,99)=18.702, *p*<0.001 for paired analysis; F(1,103)=9.130, *p*=0.003 for unpaired analysis (Fig. 2a and b). There was a significant main effect of layers using paired analysis, F(5,99)=3.700, *p*=0.004, with *post hoc* comparison indicating significantly lower sum CB1R intensity in layer VI compared to layers I and II. However, this effect was not present using unpaired analysis, F(5,103)=1.996, *p*=0.085. There was no significant condition × layer interaction in both paired and unpaired analyses, *p*=0.863 for paired; *p*=0.960 for unpaired (Fig. 2c). Significant results persisted in analyses without outlier pair (see Table S2).

**Figure 2a.**
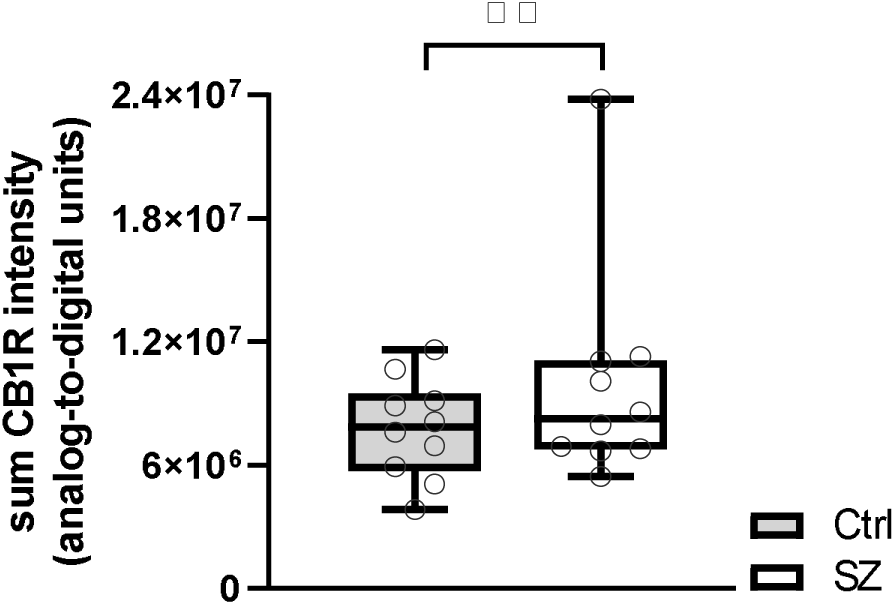
Sum CB1R intensity from postmortem PFC samples of subjects with schizophrenia (SZ) and unaffected comparisons (Ctrl). Each individual data point represents the sum intensity averaged across all sampled sites across a single subject. Central line indicates the median, box boundaries extend from the 25^th^ to 75^th^ percentiles, and whiskers extend from the minimum to maximum value. There was a main effect of subject group, *p*<0.001. \*\**p*<0.001.

**Figure 2b.**
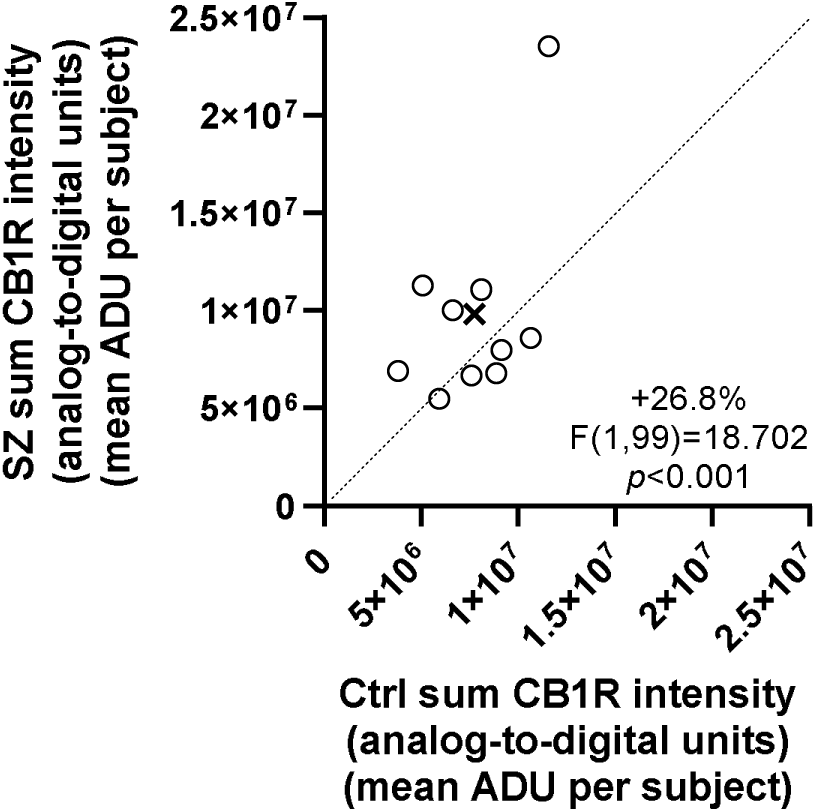
Comparison of sum CB1R intensity in matched pairs of comparison subjects (Ctrl) and subjects with schizophrenia (SZ). Mean values of sum CB1R intensities for each subject group are indicated by the X. Markers above the dashed unity line indicate pairs for which the subject with schizophrenia disorder had higher sum CB1R intensity than the matched comparison subject.

**Figure 2c.**
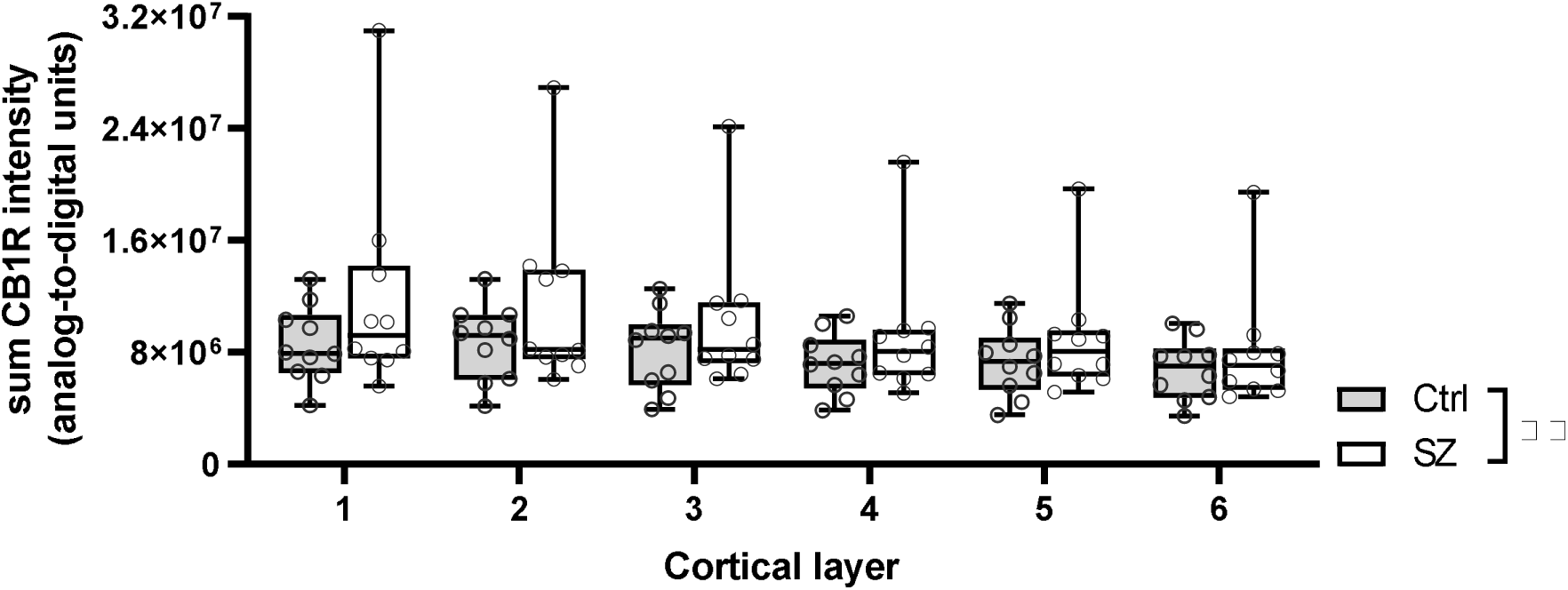
Sum CB1R intensity for individual subjects across cortical layers. Each individual data point represents the sum intensity averaged across all sampled sites for a single subject. Central line indicates the median, box boundaries extend from the 25^th^ to 75^th^ percentiles, and whiskers extend from the minimum to maximum value. There was a main effect of subject group, *p*<0.001. \*\**p*<0.001.

There were no group differences in Ctrl/SZ sum CB1R intensity ratios between pairs including SZ subjects with or without cannabis, antipsychotic, antidepressant, benzodiazepine or valproic acid exposures (Table S3).

### Terminal type-specific CB1R alterations in the PFC of patients with SZ

There was a significant main effect of terminal type in mean CB1R intensity, F(2,315)=827.566, *p*<0.001 for paired analysis; F(2,319)=786.746, *p*<0.001 for unpaired analysis; and a significant terminal type × subject group interaction, F(2,315)=22.875, *p*<0.001 for paired analysis; F(2,319)=21.747, *p*<0.001 for unpaired analysis (Fig. 3a and b). Post hoc pairwise comparisons showed 35.3% higher mean CB1R intensity in SZ compared to Ctrl within vGlut1-IR bouton populations, *p*<0.001 for both paired and unpaired analyses, and 14.9% lower mean CB1R intensity in SZ compared to Ctrl within high-CB1R-expressing vGAT-IR bouton populations, *p*<0.001 for both paired and unpaired analyses (Table 3). Significant results persisted in analyses without outlier pair (see Table S2).

**Figure 3a.**
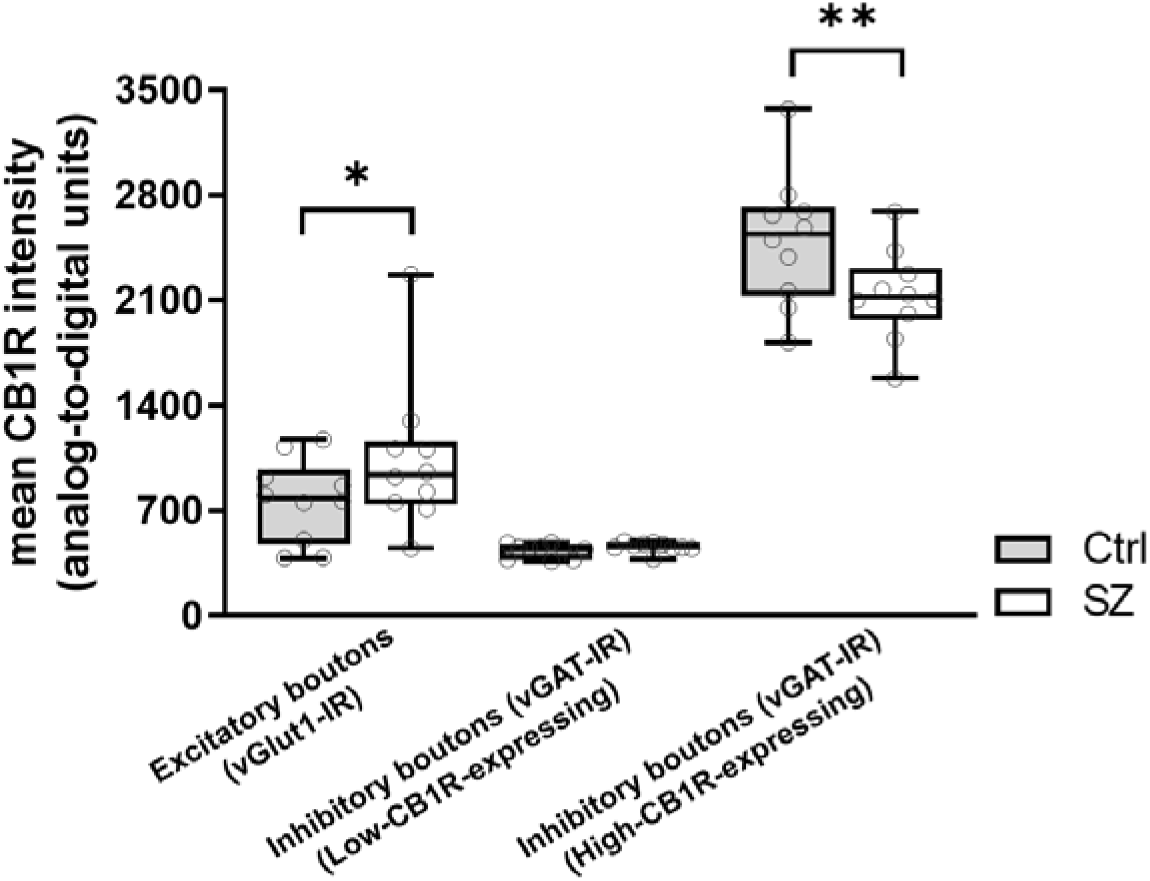
Mean CB1R intensity within excitatory (vGlut1-IR), high-CB1R-expressing inhibitory (vGAT-IR), and low-CB1R-expressing inhibitory boutons from postmortem PFC samples of subjects with schizophrenia (SZ) and unaffected comparisons (Ctrl). Each individual data point represents mean intensity averaged across all sampled sites across a single subject. Central line indicates the median, box boundaries extend from the 25^th^ to 75^th^ percentiles, and whiskers extend from the minimum to maximum value. There was a significant terminal type × subject group interaction, *p*<0.001. Mean CB1R intensity in SZ was significantly higher compared to Ctrl in excitatory boutons, and significantly lower compared to Ctrl in high-CB1R-expressing inhibitory boutons. ** *p*<0.001.

**Figure 3b.**
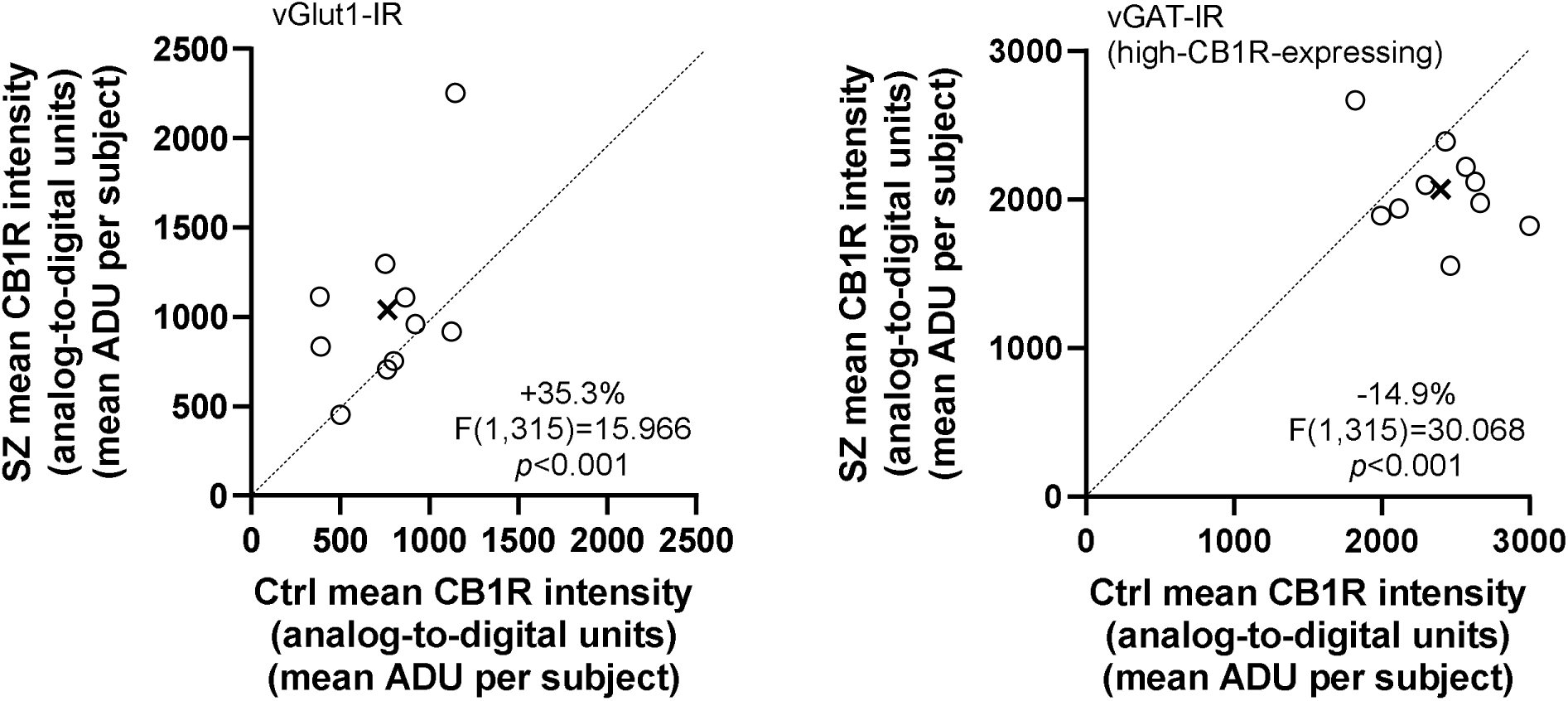
Comparison of mean CB1R intensity in excitatory (vGlut1-IR) boutons (left) and high-CB1R-expressing inhibitory (vGAT-IR) boutons (right) in matched pairs of comparison subjects (Ctrl) and subjects with schizophrenia (SZ). Mean values on mean CB1R intensities for each subject group are indicated by the X. Markers below the dashed unity line indicate pairs for which the subject with schizophrenia disorder had lower mean CB1R intensity than the matched comparison subject.

**Figure 3c.**
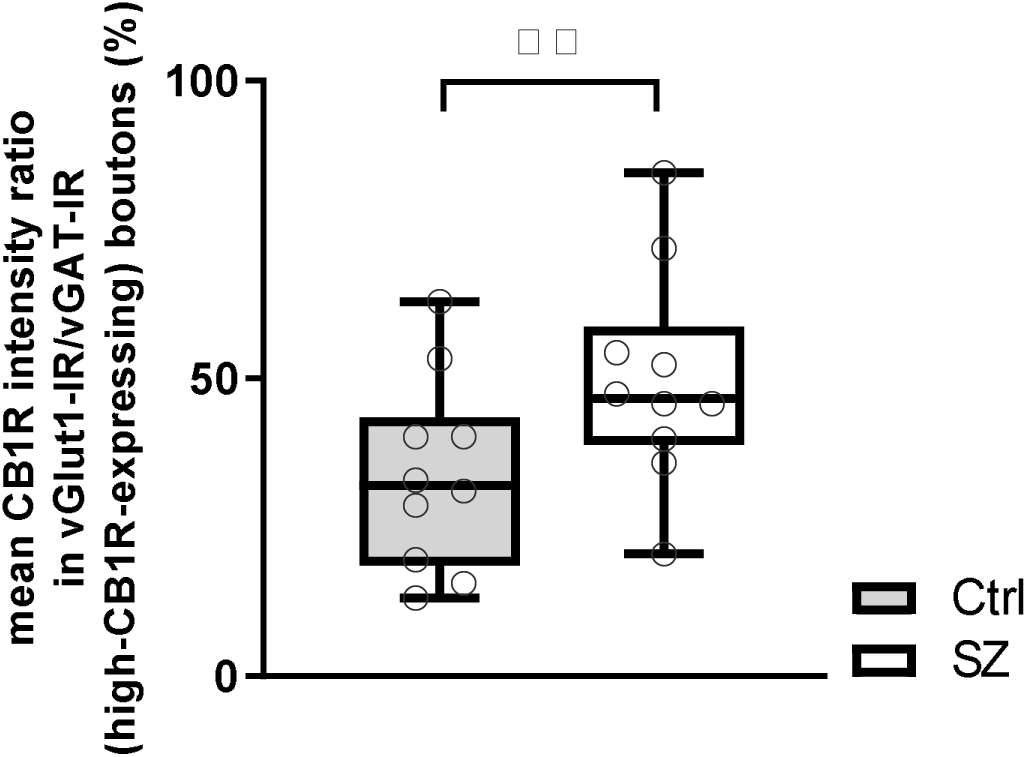
Within-pair ratios of mean CB1R intensity in vGlut-IR boutons to high-CB1R-expressing vGAT-IR boutons for individual subjects. Each individual data point represents the within-pair ratio averaged across all sampled sites for a single subject. Central line indicates the median, box boundaries extend from the 25^th^ to 75^th^ percentiles, and whiskers extend from the minimum to maximum value. There was a main effect of subject group, *p*<0.001. \*\**p*<0.001.

**Table 3.** Mean CB1R intensity in analogue-to-digital units (ADU) for all boutons across subject groups and terminal types. Values are represented as mean ± SEM.

| Terminal type | Subject group | Mean CB1R intensity (ADU ± SEM) | *p*-value |
| --- | --- | --- | --- |
| **Excitatory (vGlut1-IR)** | **Ctrl** | 770.537 ± 48.073 | *p*<0.001 |
| | **SZ** | 1042.193 ± 48.073 | |
| **High-CB1R-expressing Inhibitory (vGAT-IR)** | **Ctrl** | 2510.098 ± 48.073 | *p*<0.001 |
| | **SZ** | 2137.309 ± 48.073 | |
| **Low-CB1R-expressing Inhibitory (vGAT-IR)** | **Ctrl** | 435.461 ± 48.073 | *p*=0.714 |
| | **SZ** | 460.420 ± 48.073 | |

Abbreviations: Ctrl = unaffected comparison, SZ = schizophrenia, ADU = analog-to-digital units, SEM = standard error of mean

To further validate Terminal type-specific CB1R alterations between subject groups, we performed within subject comparisons of mean CB1R intensity ratio in vGlut1-IR to high-CB1R-expressing vGAT-IR bouton populations (Fig. 3c). There was a significant main effect of subject group, F(1,99)=53.702, *p*<0.001. The ratio of mean CB1R intensity in vGlut1-IR boutons to high-CB1R-expressing vGAT-IR boutons was 15.6% higher in SZ compared to Ctrl, indicating smaller CB1R enrichment differentials between terminal types in SZ.

There was no significant difference in mean CB1R intensity between SZ and Ctrl within the low-CB1R-expressing vGAT-IR bouton population, *p*=0.714 for paired analysis; *p*=0.802 for unpaired analysis. There was also no significant main effect of cortical layer (Figure S2; F(5,315)=1.287, *p*=0.269 for paired analysis; F(5,319)=1.223, *p*=0.298 for unpaired analysis), or significant subject group × cortical layer × terminal type three-way interaction (F(25,315)=0.758, *p*=0.794 for pair analysis; F(25,319)=0.721, *p*=0.836 for unpaired analysis).

There were no group differences in Ctrl/SZ sum CB1R intensity ratios between pairs including SZ subjects with or without cannabis, antipsychotic, antidepressant, benzodiazepine or valproic acid exposures for any terminal type (Table S4).

## Discussion

### Summary of current findings

This preliminary study compared Terminal type-specific distributions of CB1R within postmortem human PFC in individuals with SZ and non-psychiatric comparisons. We focused on this region given its involvement in the cognitive symptoms of SZ, to expand upon prior knowledge regarding CB1R alterations in this illness. When examining total CB1R, which includes not only CB1R on excitatory and inhibitory boutons, but also other locations in which CB1R are present (e.g., cholinergic, serotonergic terminals, axon segments, mitochondria) – accounting for a smaller but substantial and functionally important portion of total CB1R(62) – we identified significantly higher overall CB1R levels in individuals with SZ than non-psychiatric comparisons.

Interestingly, when examining Terminal type-specific distributions of CB1R levels, we identified a significant terminal type by subject group interaction. Specifically, mean CB1R intensity in excitatory boutons was significantly higher in SZ samples, while mean CB1R intensity in high-CB1R-expressing inhibitory boutons was significantly lower in SZ samples compared to controls.

### Comparison to prior findings

The subject pairs included in the present study were chosen based on their prior findings suggesting reciprocal alterations in CB1R protein and ligand binding. Here, our result of higher overall CB1R levels in individuals with SZ is consistent with prior results using CB1R ligand binding assays. Using postmortem brain samples, groups have assessed CB1R levels within the PFC of individuals with SZ using various radioligands, including agonist (i.e., [^3^H]CP-55940)(39,63), inverse agonist (i.e., [^3^H]MePPEP and [^3^H]-OMAR)(40,64), and antagonist (i.e., [^3^H]SR141716A)(37). Irrespective of differences in binding affinity or specificity, all studies reported higher ligand binding in samples from individuals with SZ compared to unaffected counterparts, with multiple studies controlling for covariates including age, sex, postmortem interval, THC history, and antipsychotic history. Although the same ligands as used in postmortem studies (i.e., [^11^C]MePPEP, [^11^C]OMAR) demonstrated lower global binding in individuals with SZ *in vivo*(65,66), higher global CB1R radioligand binding in SZ had similarly been demonstrated when assessed *in vivo* using the inverse agonist [^18^F]MK-9470(67). Notably, the *in vivo* studies did not specifically examine the PFC, which may account for the differing results from postmortem findings.

In addition, we expanded upon prior IHC studies examining CB1R in postmortem PFC samples from individuals with SZ, which utilized anti-CB1R antibodies that preferentially targeted high-CB1R-expressing inhibitory neurons confirmed to be CCK-positive(42). We separated mean CB1R levels in inhibitory boutons between low- and high-CB1R-expressing populations based on the median value of CB1R intensities within Ctrl samples. Our results again complemented prior findings. Specifically, we identified lower CB1R levels in postmortem PFC samples from individuals with SZ relative to comparisons when assessing the subset of high-CB1R-expressing inhibitory boutons.

### CB1R within excitatory and inhibitory neuronal populations

The current findings expand our understanding of Terminal type-specific CB1R alterations in the PFC of SZ. Here, we note that CB1R changes in SZ appear to be Terminal type-specific, with increased CB1R in excitatory terminals compared to unaffected individuals. As PFC pathology is implicated in the cognitive dysfunctions of SZ, CB1R alterations may directly contribute to symptom development by disturbing the excitatory and inhibitory balance(68) – a mechanism known to contribute to impaired salience learning(69). Considering CB1R’s role in suppressing neurotransmitter release, it is possible that these findings of higher CB1R levels in excitatory boutons of SZ represent a stronger suppression of excitatory neurotransmission (i.e., DSE). This complements the theory of glutamatergic hypofunction as a contributor to the pathology of the disorder(70,71).

Our results also identified significantly different CB1R levels in inhibitory boutons between samples from individuals with SZ and non-psychiatric comparisons, and suggested GABAergic subtype specific alterations of CB1R in SZ. Current literature supports the predominance of CB1R within CCK-containing interneurons using non-psychiatric postmortem human brain samples, with lower levels of CB1R detected in parvalbumin (PV)-positive cells using rodent studies(19). CB1R associated DSI appears to be present only within CCK-positive interneurons and not identified within other GABAergic subtypes despite low levels of CB1R being present in these interneuron populations (e.g., PV neurons)(44,60,61). Thus, our finding GABAergic CB1R alterations only within high-CB1R-expressing boutons in SZ suggest a predominant disruption of CB1R in presumptive CCK-containing interneurons, potentially contributing to the pathophysiology of the illness through attenuated DSI.

### Relationship with cannabis use in individuals with SZ

Our findings of increased CB1R in excitatory boutons and decreased CB1R in putative DSI associated inhibitory boutons in individuals with SZ may offer a potential explanation for the clinical observations of THC exposure exacerbating symptoms in SZ. An increase in CB1R within excitatory boutons may strengthen DSE following THC activation of the receptors, while a decrease in CB1R within inhibitory boutons may reduce DSI following THC exposure. It is possible that these alterations may then lead to further intensification of the glutamatergic hypoactivity present in individuals with SZ, and subsequent symptom worsening.

This is partially supported by a recent study on Terminal type-specific CB1R dependent behavioral effects using knock-out mice that underwent CB1R rescues in either dorsal telencephalic glutamatergic or forebrain GABAergic neurons(72). In CB1R knock-out mice that underwent glutamatergic CB1R rescue – a condition relevant to what we observed at present in individuals with SZ (i.e., increased glutamatergic CB1R and decreased GABAergic CB1R), THC exposure was sufficient to produce hypolocomotion. It is possible that alterations in Terminal type-specific CB1R distribution led to a disruption in E/I homeostasis, which is then exacerbated by exogenous CB1R activation through THC exposure. Additional studies using rodent manipulations would be necessary to understand how Terminal type-specific CB1R alterations may affect SZ related behaviors, and whether cannabis use leads to further Terminal type-specific behavioral disturbances under these conditions.

### Limitations

While this study expanded our understanding of CB1R alterations in SZ, its clinical generalizability is limited given its small scale and restricted subject selection. With only one pair of female subjects, three Black individuals, and predominantly middle-aged adult samples, our selection was inadequate for detecting sex, race, or age-related outcomes. However, by including these variables as covariates in our analyses, we were able to identify unique Terminal type-specific CB1R changes after controlling for these factors. Similarly, by conducting T-tests to compare results from subject pairs with or without cannabis and medication exposures, we were able to clarify that Terminal type-specific CB1R alterations observed were independent of medication or cannabis histories. However, these latter results should be interpreted in the context of limited samples. Future studies with larger sample size are needed to allow for more robust comparisons of the influence of these and other potential confounding variables.

In addition, by assessing only vGlut1 and vGAT colocalization with CB1R, the results provide only a broad overview of CB1R distributions in excitatory and inhibitory boutons, with the understanding that these groups are comprised of additional subpopulations. Future larger scale work incorporating GABAergic subtype specific markers would be necessary to fully elucidate more nuanced cell type specificity. The identity of CB1R-positive puncta not colocalized with these two markers were also unknown, and these may represent other contributors to the development of psychiatric symptoms(73).

### Conclusion and future directions

Our study replicated prior findings of higher overall CB1R levels within postmortem PFC of individuals with SZ. We also identified the presence of Terminal type-specific CB1R alterations, namely increased CB1R levels in excitatory boutons, and decreased CB1R levels in high-CB1R-expressing (presumptive CCK) inhibitory boutons in SZ. These changes suggest possible net attenuation of excitatory neurotransmission in SZ, supporting the prefrontal glutamatergic dysfunction hypothesis, lending strength to the idea that CB1R alterations disrupt PFC E/I balance in SZ. Though limitations exist, these results support the importance of conducting more in-depth CB1R examinations in SZ to elucidate the relationship between the endocannabinoid system, cannabis exposure and psychotic illnesses.

## Acknowledgments

This study was supported by MH071533 (RAS) and MH103204 (DAL).

## Supplemental information

**Table S1.**
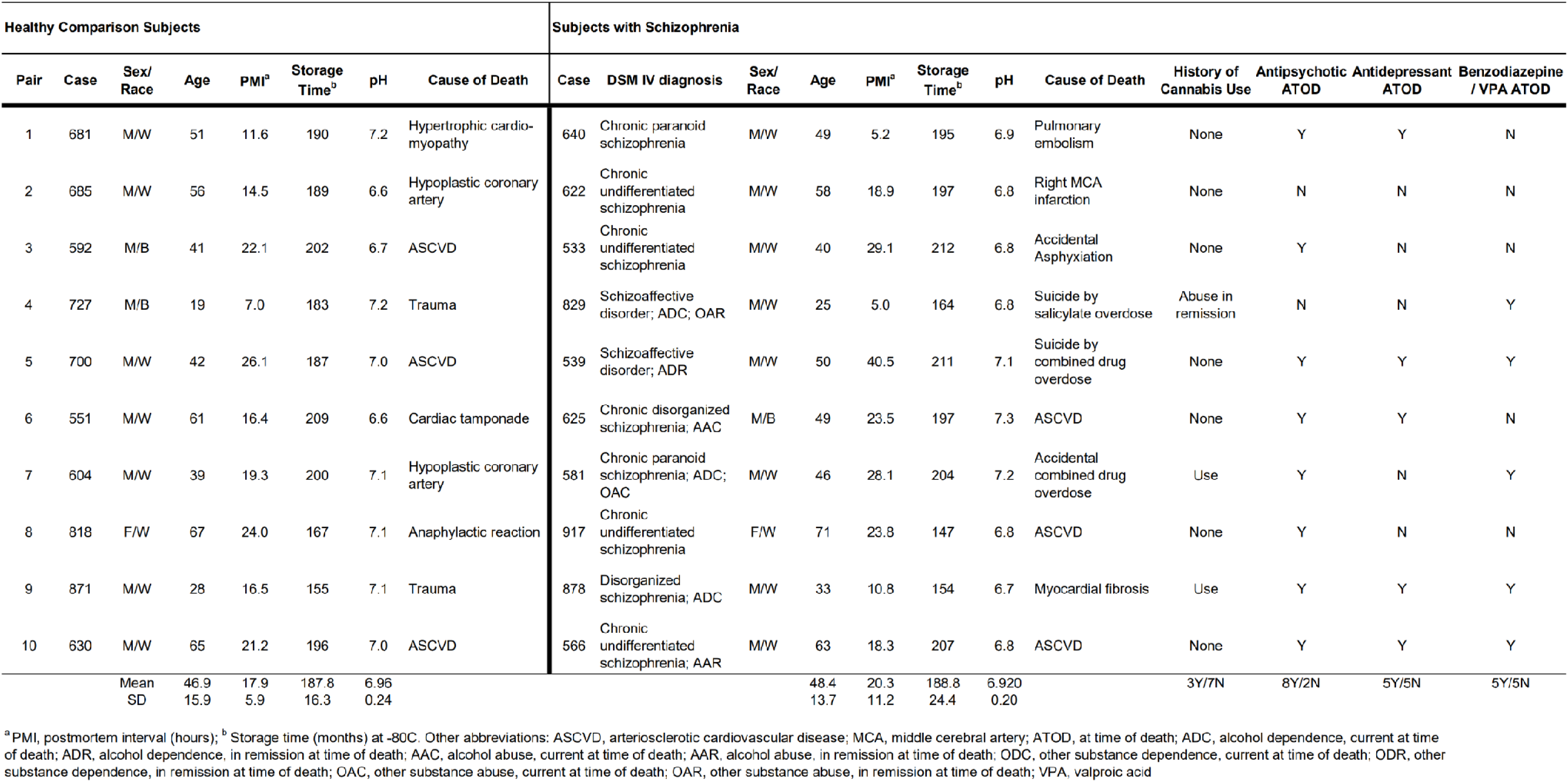
Demographic, postmortem, and clinical characteristics of individual human subjects included in the study.

**Table S2.** Statistical results of sum CB1R intensity and mean CB1R intensity analyses without outlier pair.

| Variable | Factor | p-value |
| --- | --- | --- |
| **Sum CB1R intensity** |  |  |
| **Paired analysis** | Main effect of subject group | $F_{(1,88)}=7.532, p=0.007$ |
| | Main effect of cortical layer | $F_{(5,88)}=4.803, p<0.001$ |
| **Unpaired analysis** | Main effect of subject group | $F_{(1,91)}=6.410, p=0.013$ |
| | Main effect of cortical layer | $F_{(5,91)}=4.706, p<0.001$ |
| **Mean CB1R intensity** |  |  |
| **Paired analysis** | Main effect of terminal type | $F_{(2,280)}=1053.740, p<0.001$ |
| | Subject group x terminal type interaction | $F_{(2,280)}=34.872, p<0.001$ |
|  | *Post hoc* |  |
| | vGlut-IR effect | $F_{(1,280)}=8.451, p=0.004$ |
| | high-CB1R-expressing vGAT-IR effect | $F_{(1,280)}=69.393, p<0.001$ |
| **Unpaired analysis** | Main effect of terminal type | $F_{(2,283)}=1026.559, p<0.001$ |
| | Subject group x terminal type interaction | $F_{(2,283)}=33.972, p<0.001$ |
|  | *Post hoc* |  |
| | vGlut-IR effect | $F_{(1,283)}=8.575, p=0.004$ |
| | high-CB1R-expressing vGAT-IR effect | $F_{(1,283)}=65.369, p<0.001$ |

**Table S3.** Mean values of Ctrl/SZ subject pair sum CB1R intensity ratios for pairs with and without cannabis and medication exposure history. Values are represented as mean ± SEM.

| Substance/ medication | Use history in SZ (n) | Mean of Ctrl/SZ sum CB1R intensity ratio (mean ± SEM) | *p*-value |
| --- | --- | --- | --- |
| **Cannabis** | **Yes (3)** | 95.832 ± 9.026 | t(8)=0.403 |
| | **No (7)** | 86.170 ± 4.726 | *p*=0.697 |
| **Antipsychotics** | **Yes (8)** | 91.541 ± 4.783 | t(8)=0.452 |
| | **No (2)** | 79.181 ± 9.326 | *p*=0.664 |
| **Antidepressants** | **Yes (5)** | 95.131 ± 5.866 | t(8)=0.557 |
| | **No (5)** | 83.006 ± 6.105 | *p*=0.593 |
| **Benzodiazepines/ valproic acid** | **Yes (5)** | 105.216 ± 5.977 | t(8)=1.699 |
| | **No (5)** | 72.910 ± 4.528 | *p*=0.128 |

Abbreviations: Ctrl = unaffected comparison, SZ = schizophrenia, SEM = standard error of mean

**Table S4.**
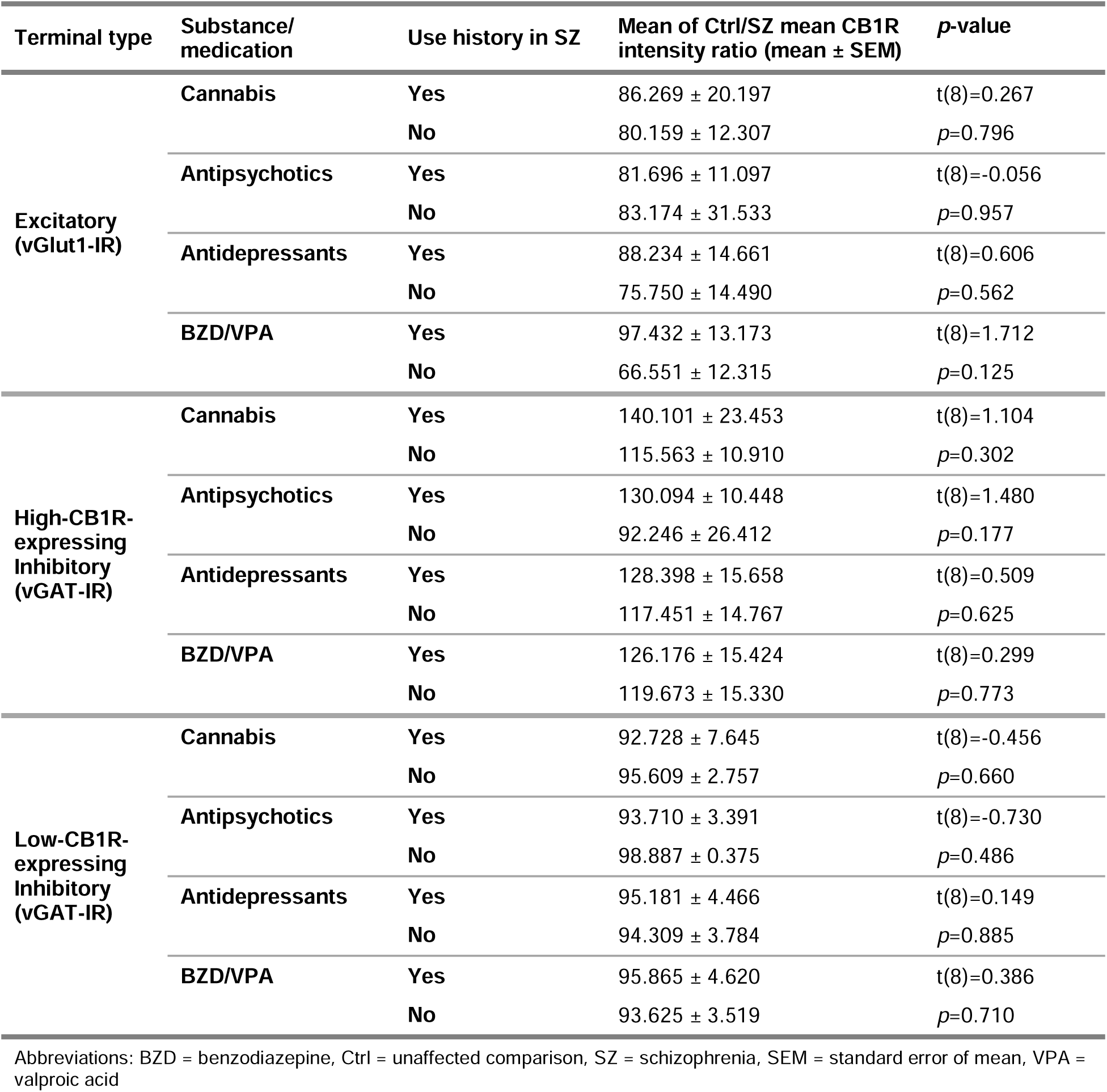
Mean values and T-tests results of Ctrl/SZ subject pair mean CB1R intensity ratios for pairs with and without cannabis and medication exposure history for each terminal type. Values are represented as mean ± SEM.

| Terminal type | Substance/<br>medication | Use history in SZ | Mean of Ctrl/SZ mean CB1R<br>intensity ratio (mean ± SEM) | p-value |
| --- | --- | --- | --- | --- |
| Excitatory<br>(vGlut1-IR) | Cannabis | Yes | 86.269 ± 20.197 | t(8)=0.267 |
| | | No | 80.159 ± 12.307 | p=0.796 |
| | Antipsychotics | Yes | 81.696 ± 11.097 | t(8)=-0.056 |
| | | No | 83.174 ± 31.533 | p=0.957 |
| | Antidepressants | Yes | 88.234 ± 14.661 | t(8)=0.606 |
| | | No | 75.750 ± 14.490 | p=0.562 |
| | BZD/VPA | Yes | 97.432 ± 13.173 | t(8)=1.712 |
| | | No | 66.551 ± 12.315 | p=0.125 |
| High-CB1R-expressing<br>Inhibitory<br>(vGAT-IR) | Cannabis | Yes | 140.101 ± 23.453 | t(8)=1.104 |
| | | No | 115.563 ± 10.910 | p=0.302 |
| | Antipsychotics | Yes | 130.094 ± 10.448 | t(8)=1.480 |
| | | No | 92.246 ± 26.412 | p=0.177 |
| | Antidepressants | Yes | 128.398 ± 15.658 | t(8)=0.509 |
| | | No | 117.451 ± 14.767 | p=0.625 |
| | BZD/VPA | Yes | 126.176 ± 15.424 | t(8)=0.299 |
| | | No | 119.673 ± 15.330 | p=0.773 |
| Low-CB1R-expressing<br>Inhibitory<br>(vGAT-IR) | Cannabis | Yes | 92.728 ± 7.645 | t(8)=-0.456 |
| | | No | 95.609 ± 2.757 | p=0.660 |
| | Antipsychotics | Yes | 93.710 ± 3.391 | t(8)=-0.730 |
| | | No | 98.887 ± 0.375 | p=0.486 |
| | Antidepressants | Yes | 95.181 ± 4.466 | t(8)=0.149 |
| | | No | 94.309 ± 3.784 | p=0.885 |
| | BZD/VPA | Yes | 95.865 ± 4.620 | t(8)=0.386 |
| | | No | 93.625 ± 3.519 | p=0.710 |

Abbreviations: BZD = benzodiazepine, Ctrl = unaffected comparison, SZ = schizophrenia, SEM = standard error of mean, VPA = valproic acid

**Figure S1.**
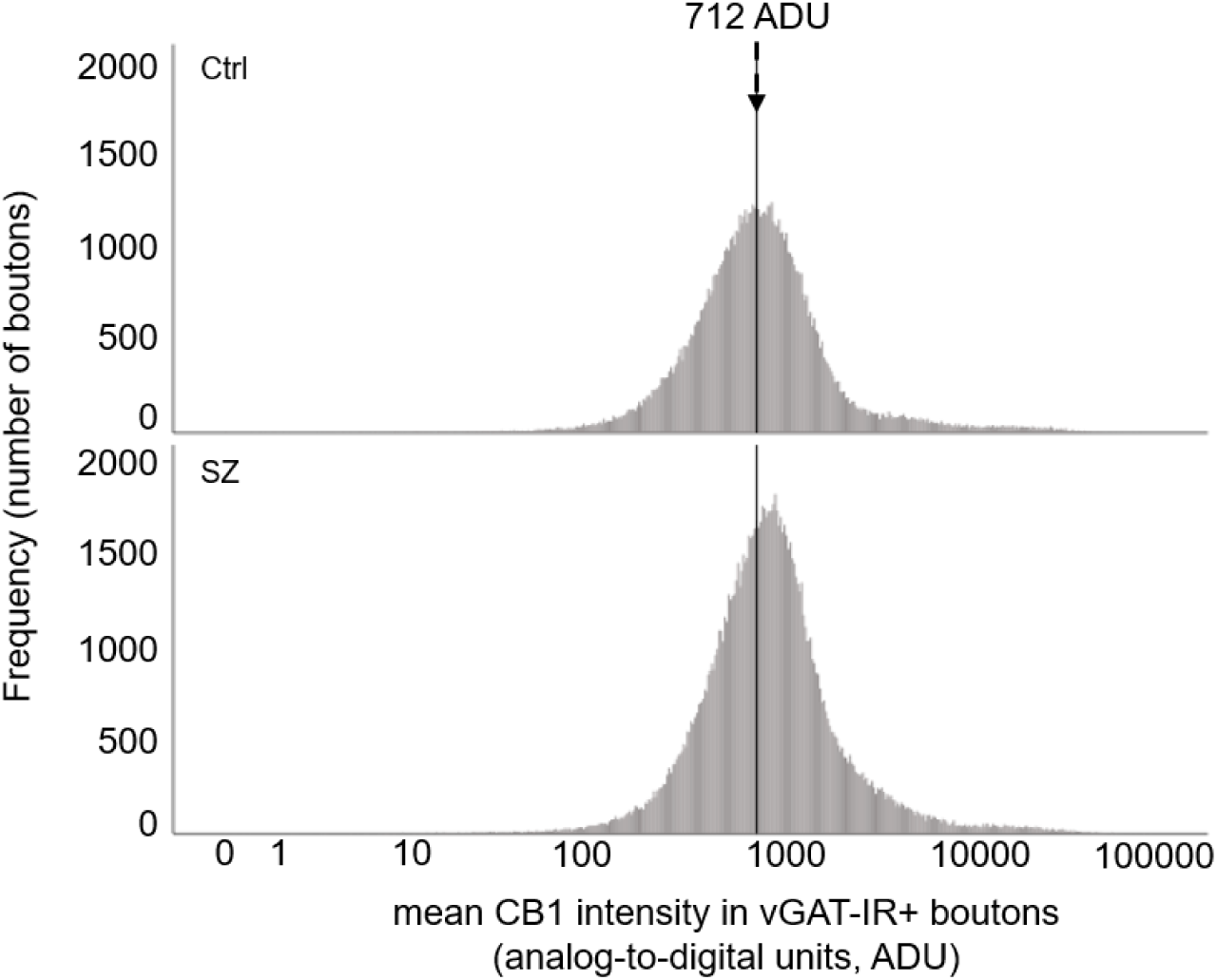
Mean CB1R intensity frequency histograms for inhibitory (vGAT-IR) boutons from postmortem PFC samples of subjects with schizophrenia (SZ) and unaffected comparisons (Ctrl), measured in analogue-to-digital units (ADU). The line at 712 ADU denotes the median value of mean CB1R intensity for vGAT-IR boutons in Ctrl.

**Figure S2.**
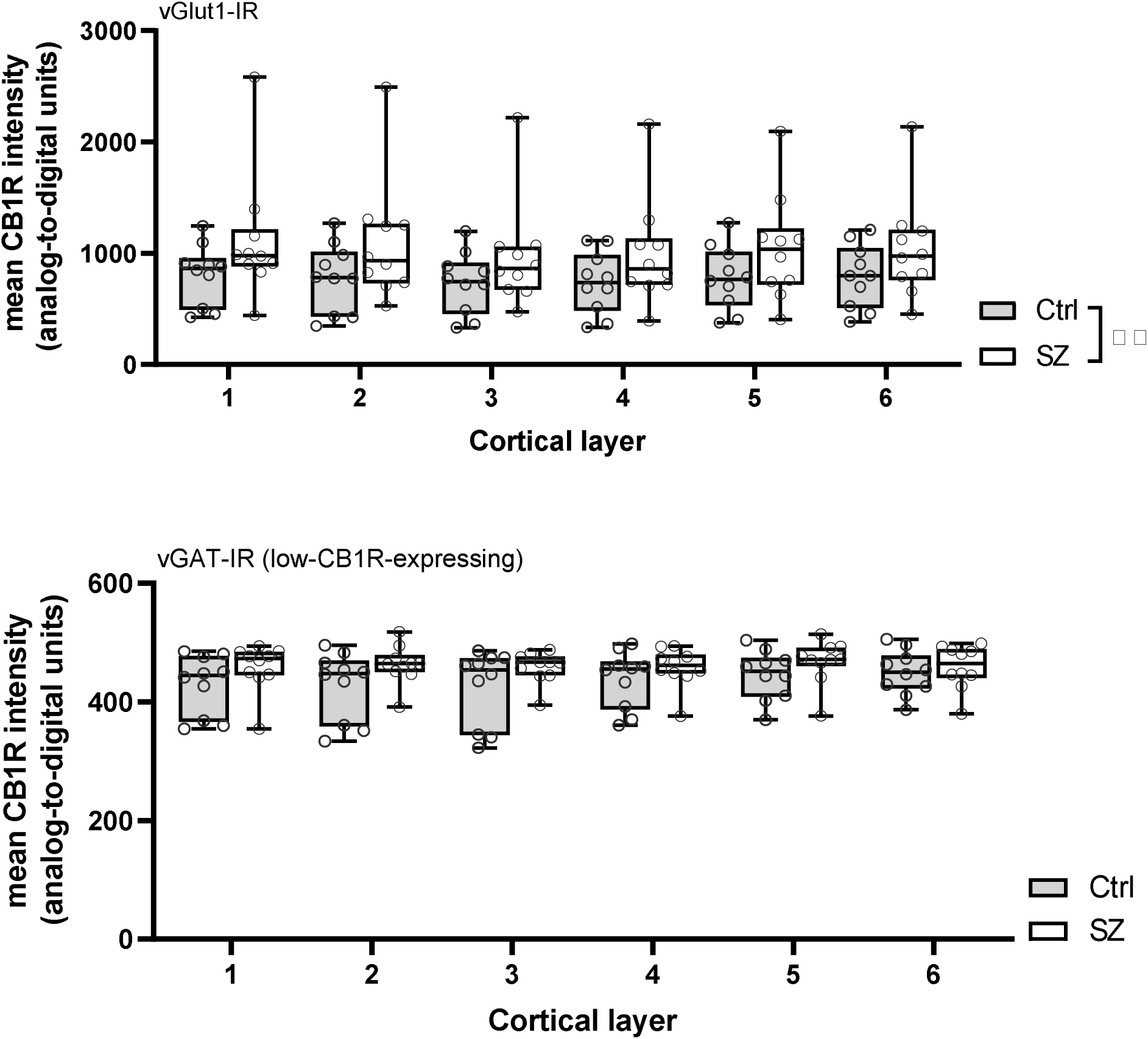
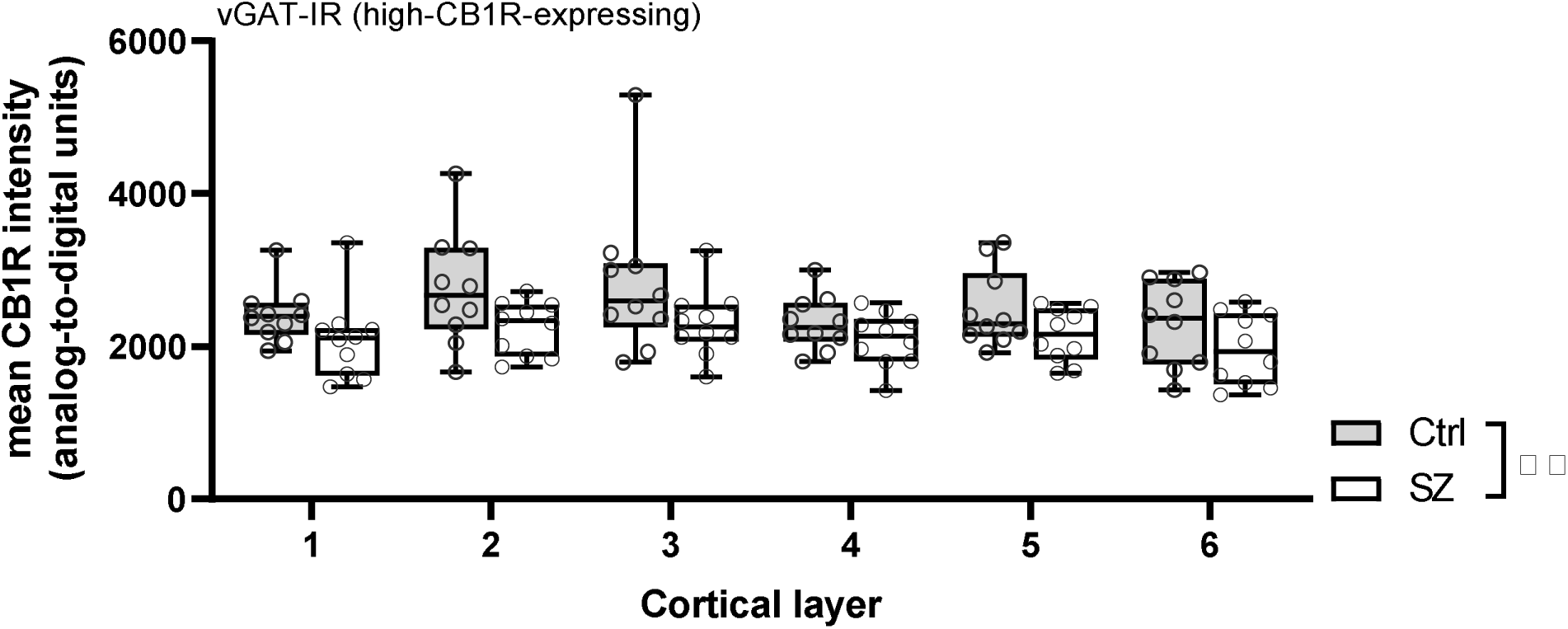
Mean CB1R intensity in excitatory (vGlut1-IR) boutons (top), low-CB1R-expressing inhibitory (vGAT-IR) boutons (middle), and high-CB1R-expressing inhibitory (vGAT-IR) boutons (bottom) for individual subjects across cortical layers. Each individual data point represents the mean intensity averaged across all sampled sites for a single subject. Central line indicates the median, box boundaries extend from the 25^th^ to 75^th^ percentiles, and whiskers extend from the minimum to maximum value. There was a main effect of subject group for excitatory and high-CB1R-expressing inhibitory boutons, *p*<0.001. \*\**p*<0.001.

## Reference

1. Hasin DS (2018, January 1): US Epidemiology of Cannabis Use and Associated Problems. Neuropsychopharmacology, vol. 43. Nature Publishing Group, pp 195–212.

2. Bhatia D, Hinckley J, Mikulich S, Sakai J (2022): Cannabis Legalization and Adolescent Use of Electronic Vapor Products, Cannabis, and Cigarettes. J Addict Med 16: E16–E22.

3. Stinson FS, Ruan WJ, Pickering R, Grant BF (2006): Cannabis use disorders in the USA: Prevalence, correlates and co-morbidity. Psychol Med 36: 1447–1460.

4. Andréasson S, Engström A, Allebeck P, Rydberg U (1987): Cannabis and schizophrenia. A Longitudinal Study of Swedish Conscripts. The Lancet 330: 1483–1486.

5. Nielsen SM, Toftdahl NG, Nordentoft M, Hjorthoj C (2017): Association between alcohol, cannabis, and other illicit substance abuse and risk of developing schizophrenia: A nationwide population based register study. Psychol Med 47: 1668–1677.

6. Callaghan RC, Cunningham JK, Allebeck P, Arenovich T, Sajeev G, Remington G, et al. (2012): Methamphetamine use and schizophrenia: A population-based cohort study in California. American Journal of Psychiatry 169: 389–396.

7. Large M, Sharma S, Compton MT, Slade T, Nielssen O (2011): Cannabis use and earlier onset of psychosis: A systematic meta-analysis. Arch Gen Psychiatry 68: 555–561.

8. Zammit S, Allebeck P, Andreasson S, Lundberg I, Lewis G (2002): Self reported cannabis use as a risk factor for schizophrenia in Swedish conscripts of 1969: Historical cohort study. Br Med J 325: 1199– 1201.

9. Arendt M, Rosenberg R, Foldager L, Perto G, Munk-Jørgensen P (2005): Cannabis-induced psychosis and subsequent schizophrenia-spectrum disorders: Follow-up study of 535 incident cases. British Journal of Psychiatry 187: 510–515.

10. Henquet C, Krabbendam L, Spauwen J, Kaplan C, Lieb R, Wittchen HU, van Os J (2005): Prospective cohort study of cannabis use, predisposition for psychosis, and psychotic symptoms in young people. Br Med J 330: 11–14.

11. D’Souza DC, Abi-Saab WM, Madonick S, Forselius-Bielen K, Doersch A, Braley G, et al. (2005): Delta-9-tetrahydrocannabinol effects in schizophrenia: Implications for cognition, psychosis, and addiction. Biol Psychiatry 57: 594–608.

12. Manrique-Garcia E, Zammit S, Dalman C, Hemmingsson T, Andreasson S, Allebeck P (2014): Prognosis of schizophrenia in persons with and without a history of cannabis use. Psychol Med 44: 2513–2521.

13. Patel SJ, Khan S, M S, Hamid P (2020): The Association Between Cannabis Use and Schizophrenia: Causative or Curative? A Systematic Review. Cureus 12. https://doi.org/10.7759/CUREUS.9309

14. Gage SH, Jones HJ, Burgess S, Bowden J, Davey Smith G, Zammit S, Munafò MR (2017): Assessing causality in associations between cannabis use and schizophrenia risk: A two-sample Mendelian randomization study. Psychol Med 47: 971–980.

15. Vaucher J, Keating BJ, Lasserre AM, Gan W, Lyall DM, Ward J, et al. (2018): Cannabis use and risk of schizophrenia: A Mendelian randomization study. Mol Psychiatry 23: 1287–1292.

16. Shrivastava A, Johnston M, Terpstra K, Bureau Y (2014): Cannabis and psychosis: Neurobiology. Indian J Psychiatry 56: 8.

17. Busquets-Garcia A, Bains J, Marsicano G (2018, January 1): CB 1 Receptor Signaling in the Brain: Extracting Specificity from Ubiquity. Neuropsychopharmacology, vol. 43. Nature Publishing Group, pp 4–20.

18. Trettel J, Fortin DA, Levine ES (2004): Endocannabinoid signalling selectively targets perisomatic inhibitory inputs to pyramidal neurones in juvenile mouse neocortex. Journal of Physiology 556: 95–107.

19. Katona I, Sperlágh B, Sík A, Käfalvi A, Vizi ES, Mackie K, Freund TF (1999): Presynaptically located CB1 cannabinoid receptors regulate GABA release from axon terminals of specific hippocampal interneurons. Journal of Neuroscience 19: 4544–4558.

20. Bellocchio L, Lafentre P, Cannich A, Cota D, Puente N, Grandes P, et al. (2010): Bimodal control of stimulated food intake by the endocannabinoid system. Nat Neurosci 13: 281–283.

21. Chou S, Ranganath T, Fish KN, Lewis DA, Sweet RA (2022): Cell type specific cannabinoid CB1 receptor distribution across the human and non-human primate cortex. Sci Rep 12: 9605.

22. Egertová M, Elphick MR (2000): Localisation of cannabinoid receptors in the rat brain using antibodies to the intracellular C-terminal tail of CB. J Comp Neurol 422: 159–171.

23. Trettel J, Levine ES (2002): Cannabinoids depress inhibitory synaptic inputs received by layer 2/3 pyramidal neurons of the neocortex. J Neurophysiol 88: 534–539.

24. Fortin DA, Levine ES (2007): Differential effects of endocannabinoids on glutamatergic and GABAergic inputs to layer 5 pyramidal neurons. Cerebral Cortex 17: 163–174.

25. den Boon FS, Werkman TR, Schaafsma-Zhao Q, Houthuijs K, Vitalis T, Kruse CG, et al. (2015): Activation of type-1 cannabinoid receptor shifts the balance between excitation and inhibition towards excitation in layer II/III pyramidal neurons of the rat prelimbic cortex. Pflugers Arch 467: 1551–1564.

26. Guo JY, Ragland JD, Carter CS (2019, May 1): Memory and cognition in schizophrenia. Molecular Psychiatry, vol. 24. Nature Publishing Group, pp 633–642.

27. Garey LJ, Ong Y, Patel TS, Kanani M, Davis A, Mortimer AM, et al. (1998): Reduced dendritic spine density on cerebral cortical pyramidal neurons in schizophrenia. J Neurol Neurosurg Psychiatry 65: 446–453.

28. Konopaske GT, Lange N, Coyle JT, Benes FM (2014): Prefrontal Cortical Dendritic Spine Pathology in Schizophrenia and Bipolar Disorder. JAMA Psychiatry 71: 1323–1331.

29. Glantz LA, Lewis DA (2000): Decreased dendritic spine density on prefrontal cortical pyramidal neurons in schizophrenia. Arch Gen Psychiatry 57: 65–73.

30. Smucny J, Dienel SJ, Lewis DA, Carter CS (n.d.): Mechanisms underlying dorsolateral prefrontal cortex contributions to cognitive dysfunction in schizophrenia. https://doi.org/10.1038/s41386-021-01089-0

31. Glausier JR, Enwright JF, Lewis DA (2020): Diagnosis- and Cell Type-Specific Mitochondrial Functional Pathway Signatures in Schizophrenia and Bipolar Disorder. Am J Psychiatry 177: 1140–1150.

32. Hashimoto T, Volk DW, Eggan SM, Mirnics K, Pierri JN, Sun Z, et al. (2003): Gene expression deficits in a subclass of GABA neurons in the prefrontal cortex of subjects with schizophrenia. Journal of Neuroscience 23: 6315–6326.

33. Dienel SJ, Lewis DA (2019): Alterations in cortical interneurons and cognitive function in schizophrenia. Neurobiol Dis 131. https://doi.org/10.1016/J.NBD.2018.06.020

34. Ferretjans R, Moreira FA, Teixeira AL, Salgado J v. (2012): The endocannabinoid system and its role in schizophrenia: A systematic review of the literature. Revista Brasileira de Psiquiatria 34: 163–193.

35. Ibarra-Lecue I, Pilar-Cuéllar F, Muguruza C, Florensa-Zanuy E, Díaz Á, Urigüen L, et al. (2018, November 1): The endocannabinoid system in mental disorders: Evidence from human brain studies. Biochemical Pharmacology, vol. 157. Elsevier Inc., pp 97–107.

36. Dean B, Sundram S, Bradbury R, Scarr E, Copolov DD (2001): Studies on [3H]CP-55940 binding in the human central nervous system: Regional specific changes in density of cannabinoid-1 receptors associated with schizophrenia and cannabis use. Neuroscience 103: 9–15.

37. Zavitsanou K, Garrick T, Huang XF (2004): Selective antagonist [3H]SR141716A binding to cannabinoid CB1 receptors is increased in the anterior cingulate cortex in schizophrenia. Prog Neuropsychopharmacol Biol Psychiatry 28: 355–360.

38. Newell KA, Deng C, Huang XF (2006): Increased cannabinoid receptor density in the posterior cingulate cortex in schizophrenia. Exp Brain Res 172: 556–560.

39. Dalton VS, Long LE, Weickert CS, Zavitsanou K (2011): Paranoid schizophrenia is characterized by increased CB 1 receptor binding in the dorsolateral prefrontal cortex. Neuropsychopharmacology 36: 1620–1630.

40. Volk DW, Eggan SM, Horti AG, Wong DF, Lewis DA (2014): Reciprocal alterations in cortical cannabinoid receptor 1 binding relative to protein immunoreactivity and transcript levels in schizophrenia. Schizophr Res 159: 124–129.

41. Urigüen L, García-Fuster MJ, Callado LF, Morentin B, la Harpe R, Casadó V, et al. (2009): Immunodensity and mRNA expression of A2A adenosine, D 2 dopamine, and CB1 cannabinoid receptors in postmortem frontal cortex of subjects with schizophrenia: Effect of antipsychotic treatment. Psychopharmacology (Berl) 206: 313–324.

42. Eggan SM, Stoyak SR, Verrico CD, Lewis DA (2010): Cannabinoid CB1 Receptor Immunoreactivity in the Prefrontal Cortex: Comparison of Schizophrenia and Major Depressive Disorder. Neuropsychopharmacology 35: 2060–2071.

43. Eggan SM, Hashimoto T, Lewis DA (2008): Reduced cortical cannabinoid 1 receptor messenger RNA and protein expression in schizophrenia. Arch Gen Psychiatry 65: 772–784.

44. Gonzalez-Burgos G, Miyamae T, Pafundo DE, Yoshino H, Rotaru DC, Hoftman G, et al. (2015): Functional Maturation of GABA Synapses During Postnatal Development of the Monkey Dorsolateral Prefrontal Cortex. Cerebral Cortex 25: 4076–4093.

45. DeGiosio R, Kelly RM, DeDionisio AM, Newman JT, Fish KN, Sampson AR, et al. (2019): MAP2 immunoreactivity deficit is conserved across the cerebral cortex within individuals with schizophrenia. NPJ Schizophr 5. https://doi.org/10.1038/s41537-019-0081-0

46. Jiao Y, Sun Z, Lee T, Fusco FR, Kimble TD, Meade CA, et al. (1999): A simple and sensitive antigen retrieval method for free-floating and slide-mounted tissue sections. J Neurosci Methods 93: 149– 162.

47. Clancy B, Cauller LJ (1998): Reduction of background autofluorescence in brain sections following immersion in sodium borohydride. J Neurosci Methods 83: 97–102.

48. Rocco BR, DeDionisio AM, Lewis DA, Fish KN (2017): Alterations in a Unique Class of Cortical Chandelier Cell Axon Cartridges in Schizophrenia. Biol Psychiatry 82: 40–48.

49. Moyer CE, Delevich KM, Fish KN, Asafu-Adjei JK, Sampson AR, Dorph-Petersen KA, et al. (2013): Intracortical excitatory and thalamocortical boutons are intact in primary auditory cortex in schizophrenia. Schizophr Res 149: 127–134.

50. Rocco BR, DeDionisio AM, Lewis DA, Fish KN (2017): Alterations in a unique class of cortical chandelier cell axon cartridges in schizophrenia. Biol Psychiatry 82: 40.

51. Rocco BR, Sweet RA, Lewis DA, Fish KN (2016): GABA-Synthesizing Enzymes in Calbindin and Calretinin Neurons in Monkey Prefrontal Cortex. Cerebral Cortex (New York, NY) 26: 2191.

52. Morini R, Ghirardini E, Butti E, Verderio C, Martino G, Matteoli M (2015): Subventricular zone neural progenitors reverse TNF-alpha effects in cortical neurons. Stem Cell Res Ther 6. https://doi.org/10.1186/s13287-015-0158-2

53. Martens H, Weston MC, Boulland JL, Grønborg M, Grosche J, Kacza J, et al. (2008): Unique luminal localization of VGAT-C terminus allows for selective labeling of active cortical GABAergic synapses. J Neurosci 28: 13125–13131.

54. Barnes JL, Mohr C, Ritchey CR, Erikson CM, Shiina H, Rossi DJ (2020): Developmentally Transient CB1Rs on Cerebellar Afferents Suppress Afferent Input, Downstream Synaptic Excitation, and Signaling to Migrating Neurons. J Neurosci 40: 6133–6145.

55. Sweet RA, Fish KN, Lewis DA (2010): Mapping synaptic pathology within cerebral cortical circuits in subjects with schizophrenia. Front Hum Neurosci 4: 1–14.

56. DeGiosio R, Kelly RM, DeDionisio AM, Newman JT, Fish KN, Sampson AR, et al. (2019): MAP2 immunoreactivity deficit is conserved across the cerebral cortex within individuals with schizophrenia. NPJ Schizophr 5: 1–9.

57. Pierri JN, Chaudry AS, Woo T-UW, Lewis DA (1999): Alterations in Chandelier Neuron Axon Terminals in the Prefrontal Cortex of Schizophrenic Subjects. Am J Psychiatry 156: 11.

58. McKinney BC, MacDonald ML, Newman JT, Shelton MA, DeGiosio RA, Kelly RM, et al. (2019): Density of small dendritic spines and microtubule-associated-protein-2 immunoreactivity in the primary auditory cortex of subjects with schizophrenia. Neuropsychopharmacology 44: 1055–1061.

59. Fish KN, Sweet RA, Deo AJ, Lewis DA (2008): An automated segmentation methodology for quantifying immunoreactive puncta number and fluorescence intensity in tissue sections. Brain Res 1240: 62–72.

60. Karson MA, Whittington KC, Alger BE (2008): Cholecystokinin inhibits endocannabinoid-sensitive hippocampal IPSPs and stimulates others. Neuropharmacology 54: 117–128.

61. Iball J, Ali AB (2011): Endocannabinoid Release Modulates Electrical Coupling between CCK Cells Connected via Chemical and Electrical Synapses in CA1. Front Neural Circuits 5. https://doi.org/10.3389/FNCIR.2011.00017

62. Lu H-C, Mackie K (2021): Review of the Endocannabinoid System. Biol Psychiatry Cogn Neurosci Neuroimaging 6: 607–615.

63. Dean B, Sundram S, Bradbury R, Scarr E, Copolov DD (2001): Studies on [3H]CP-55940 binding in the human central nervous system: Regional specific changes in density of cannabinoid-1 receptors associated with schizophrenia and cannabis use. Neuroscience 103: 9–15.

64. Jenko KJ, Hirvonen J, Henter ID, Anderson KB, Zoghbi SS, Hyde TM, et al. (2012): Binding of a tritiated inverse agonist to cannabinoid CB1 receptors is increased in patients with schizophrenia. Schizophr Res 141: 185–188.

65. Borgan F, Laurikainen H, Veronese M, Marques TR, Haaparanta-Solin M, Solin O, et al. (2019): In Vivo Availability of Cannabinoid 1 Receptor Levels in Patients with First-Episode Psychosis. JAMA Psychiatry 76: 1074–1084.

66. Ranganathan M, Cortes-Briones J, Radhakrishnan R, Thurnauer H, Planeta B, Skosnik P, et al. (2016): Reduced Brain Cannabinoid Receptor Availability in Schizophrenia. Biol Psychiatry 79: 997–1005.

67. Ceccarini J, de Hert M, van Winkel R, Peuskens J, Bormans G, Kranaster L, et al. (2013): Increased ventral striatal CB1 receptor binding is related to negative symptoms in drug-free patients with schizophrenia. Neuroimage 79: 304–312.

68. den Boon FS, Werkman TR, Schaafsma-Zhao Q, Houthuijs K, Vitalis T, Kruse CG, et al. (2015): Activation of type-1 cannabinoid receptor shifts the balance between excitation and inhibition towards excitation in layer II/III pyramidal neurons of the rat prelimbic cortex. Pflugers Arch 467: 1551–1564.

69. Insel N, Guerguiev J, Richards BA (2018): Irrelevance by inhibition: Learning, computation, and implications for schizophrenia. PLoS Comput Biol 14. https://doi.org/10.1371/journal.pcbi.1006315

70. Dixon BJ, Kumar J, Danielmeier C (2022): Frontal neural metabolite changes in schizophrenia and their association with cognitive control: A systematic review. Neurosci Biobehav Rev 132: 224–247.

71. Dauvermann MR, Lee G, Dawson N (2017): Glutamatergic regulation of cognition and functional brain connectivity: insights from pharmacological, genetic and translational schizophrenia research. Br J Pharmacol 174: 3136–3160.

72. de Giacomo V, Ruehle S, Lutz B, Häring M, Remmers F (2020): Differential glutamatergic and GABAergic contributions to the tetrad effects of Δ 9-tetrahydrocannabinol revealed by cell-type-specific reconstitution of the CB1 receptor. Neuropharmacology 179. https://doi.org/10.1016/J.NEUROPHARM.2020.108287

73. Han X, He Y, Bi GH, Zhang HY, Song R, Liu QR, et al. (2017): CB1 Receptor Activation on VgluT2-Expressing Glutamatergic Neurons Underlies Δ 9-Tetrahydrocannabinol (Δ 9-THC)-Induced Aversive Effects in Mice. Sci Rep 7. https://doi.org/10.1038/S41598-017-12399-Z

